# Stimulation Artifact Source Separation (SASS) for assessing electric brain oscillations during transcranial alternating current stimulation (tACS)

**DOI:** 10.1101/2020.04.03.023192

**Authors:** David Haslacher, Khaled Nasr, Stephen E. Robinson, Christoph Braun, Surjo R. Soekadar

## Abstract

Brain oscillations, e.g. measured by electro- or magnetoencephalography (EEG/MEG), are causally linked to brain functions that are fundamental for perception, cognition and learning. Recent advances in neurotechnology provide means to non-invasively target these oscillations using frequency-tuned amplitude-modulated transcranial alternating current stimulation (AM-tACS). However, online adaptation of stimulation parameters to ongoing brain oscillations remains an unsolved problem due to stimulation artifacts that impede such adaptation, particularly at the target frequency. Here, we introduce a real-time compatible artifact rejection algorithm (Stimulation Artifact Source Separation, SASS) that overcomes this limitation. SASS is a spatial filter (linear projection) removing EEG signal components that are maximally different in the presence versus absence of stimulation. This enables the reliable removal of stimulation-specific signal components, while leaving physiological signal components unaffected. For validation of SASS, we evoked brain activity with known phase and amplitude using 10 Hz visual flickers across 7 healthy human volunteers. 64-channel EEG was recorded during and in absence of 10 Hz AM-tACS targeting the visual cortex. Phase differences between AM-tACS and the visual stimuli were randomized, so that steady-state visually evoked potentials (SSVEPs) were phase-locked to the visual stimuli but not to the AM-tACS signal. For validation, distributions of single-trial amplitude and phase of EEG signals recorded during and in absence of AM-tACS were compared for each participant. When no artifact rejection method was applied, AM-tACS stimulation artifacts impeded assessment of single-trial SSVEP amplitude and phase. Using SASS, amplitude and phase of single trials recorded during and in absence of AM-tACS were comparable. These results indicate that SASS can be used to establish adaptive (closed-loop) AM-tACS, a potentially powerful tool to target various brain functions, and to investigate how AM-tACS interacts with electric brain oscillations.

**Highlights:**

- Stimulation Artifact Source Separation (SASS), a real-time compatible signal decomposition algorithm for separating electric brain activity and stimulation signal artifacts related to amplitude-modulated transcranial alternating current stimulation (AM-tACS), is introduced
- Employing SASS, phase and amplitude of single-trial steady state visual evoked potentials (SSVEPs) were reliably recovered from electroencephalography (EEG) recordings at the frequency targeted with AM-tACS
- SASS enables assessment of single-trial oscillatory brain activity at the target frequency during stimulation and paves the way for online adaptation of stimulation parameters to ongoing brain oscillations

## 1. Introduction

Brain oscillations reflect neuronal cell-assembly formation causally linked to various brain functions, such as perception (Fries, Schroder, Roelfsema, Singer, & Engel, 2002; Hipp, Engel, & Siegel, 2011; Rodriguez et al., 1999), cognition (Kahana, Sekuler, Caplan, Kirschen, & Madsen, 1999), memory (Fell et al., 2001) and learning (Miltner, Braun, Arnold, Witte, & Taub, 1999; Seager, Johnson, Chabot, Asaka, & Berry, 2002). While building on fine-tuned neurochemical processes at the cellular level, brain oscillations were found to be closely related to cortico-cortical communication at the neural circuit and system level (Kopell, Ermentrout, Whittington, & Traub, 2000; Roelfsema, Engel, Konig, & Singer, 1997). As such, brain oscillations assessed by electro- or magnetoencephalography (EEG/MEG) may represent a valuable target to treat neurological and psychiatric disorders in which phase synchronization and large-scale integration is disturbed, e.g. Parkinson’s disease, depression or schizophrenia.

A well-established tool to non-invasively target oscillatory brain activity uses transcranial alternating currents specifically tuned to physiological frequencies, e.g. in the alpha (8-15 Hz) or beta band (15-30 Hz). When targeting such frequencies, distinct effects on perception (Helfrich et al., 2014; Thut et al., 2017), movement (Wach et al., 2013), memory (Reinhart & Nguyen, 2019) or emotion regulation (Clancy et al., 2018) were demonstrated. While very promising in its application (Helfrich et al., 2014; Thut, Schyns, & Gross, 2011), tuning of stimulation parameters to ongoing brain oscillations (e.g. frequency, phase, intensity, and spatial distribution of the electric fields using multi-electrode montages) was unfeasible up to now because stimulation artifacts impede reliable reconstruction of physiological brain activity. Typically, the largest tACS artifact appears at the target frequency at which energy density is highest. A complicating issue relates to nonlinear modulations of this artifact by non-physiological (such as hardware- or signal processing-related) and physiological processes (such as heartbeat and respiration) (Noury, Hipp, & Siegel, 2016; Noury & Siegel, 2017b). These result in additional stimulation artifacts at other frequencies including second- and higher-order intermodulation distortions. Despite the development of a variety of tACS artifact suppression strategies such as template subtraction (Helfrich et al., 2014), adaptive filtering (Kohli & Casson, 2019), spatial filtering using beamforming (Neuling, Ruhnau, Weisz, Herrmann, & Demarchi, 2017) or signal-space projection (Vosskuhl, Mutanen, Neuling, Ilmoniemi, & Herrmann, 2019), there is currently no established artifact suppression strategy available that allows for real-time tuning of tACS stimulation parameters to the targeted brain oscillation. This not only limits effective targeting of ongoing brain oscillations, but also the possibility to systematically investigate how tACS interacts with endogenous rhythmic brain activity, a critical prerequisite to develop new and effective closed-loop adaptive brain stimulation protocols.

Recently, we have introduced a novel tACS approach that uses amplitude modulation of a high frequency carrier signal (e.g. 220 Hz) to reduce stimulation-related artifacts at the lower physiological frequency bands (Witkowski et al., 2016). By modulating the carrier signal’s amplitude at a physiological frequency (target frequency), specific brain functions could be influenced, e.g. working memory performance when targeting frontal midline theta (FMT) oscillations (Chander et al., 2016). This finding was corroborated by computational simulations showing that AM-tACS leads to phase-locking of cortical oscillations with the stimulation signal and suggests that AM-tACS exhibits the same target engagement mechanism as conventional (unmodulated) low frequency tACS (Negahbani, Kasten, Herrmann, & Fröhlich, 2018), although possibly requiring higher current density. While AM-tACS can substantially reduce stimulation artifact contamination of the targeted physiological frequency band, similar to conventional tACS, nonlinearities related to non-physiological and physiological processes (Noury et al., 2016; Noury & Siegel, 2017b) as well as intermodulation distortions could be mistaken for neural entrainment (Kasten, Negahbani, Fröhlich, & Herrmann, 2018). It would be thus important to establish an approach that efficiently separates endogenous brain activity at the target frequency from signal components related to AM-tACS artifacts.

Here, we introduce Stimulation Artifact Source Separation (SASS), a real-time-compatible signal decomposition algorithm, that allows for separating electric brain activity and AM-tACS stimulation artifacts. To test validity and reliability of SASS, brain oscillations with known amplitude and phase were evoked using a 10 Hz steady-state visual evoked potential (SSVEP) paradigm. Brain oscillations were recorded by 64-channel EEG while 10-Hz AM-tACS was applied over the visual cortex. In each trial, phase of AM-tACS relative to the visual flicker was randomly selected, such that the artifact eliminated the consistent phase relationship (phase locking) between the flicker and EEG signal. Using SASS, we successfully recovered single-trial phase and amplitude information of SSVEPs across six healthy human volunteers.

## 2. Material and Methods

### 2.1 Stimulation Artifact Source Separation (SASS)

#### 2.1.1 Overview

SASS is a spatial filter applied to encephalographic data contaminated by transcranial electric stimulation (tES) artifacts. SASS is computed from two covariance matrices, obtained from data bandpass filtered around the stimulation frequency during AM-tACS and in absence of AM-tACS. SASS is computed from and applied to the full-length recordings before any further analysis. It should be noted that, depending on the signal of interest, the SASS projection matrix **P** can be applied to the filtered or unfiltered data. In the present work, we apply **P** to the filtered data for single-trial SSVEP analysis as well as to broadband data for power spectral analysis. The entire procedure is outlined in Figure 1.

**Figure 1:**
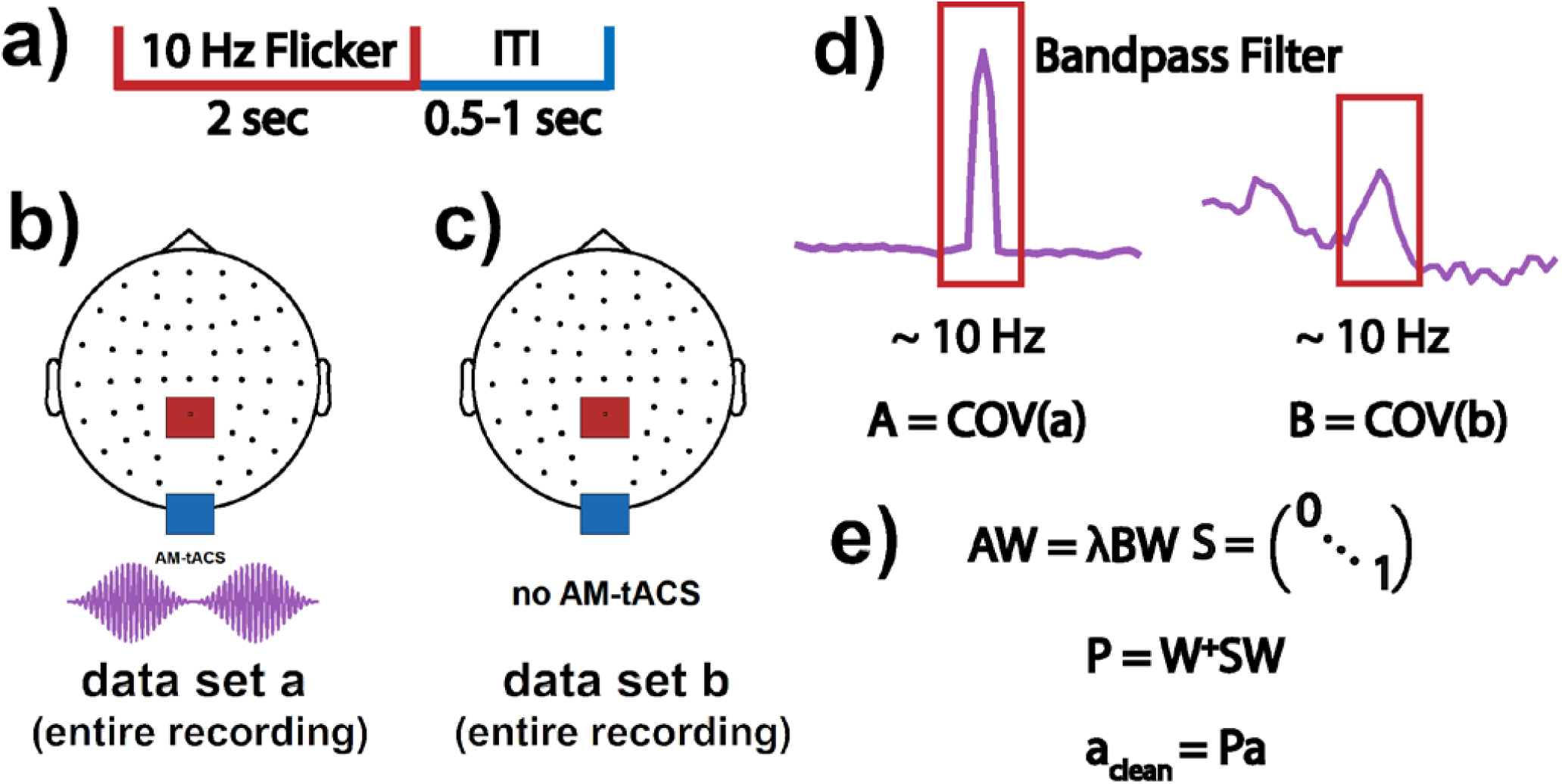
Depiction of applying Stimulation Artifact Source Separation (SASS) to electroencephalographic (EEG) data. Initially, EEG data is recorded during the paradigm of interest (A) (e.g., a 10-Hz steady-state visually evoked potentials, SSVEP, paradigm) while amplitude-modulated transcranial alternating current stimulation (AM-tACS) is applied (B) or not applied (C). These full-length recordings are then bandpass filtered around the target frequency (i.e., 10 Hz), and their narrowband covariance matrices A and B are computed (D). Finally, the SASS projection matrix P is computed from a joint diagonalization of A and B and applied to the data before any further processing (E).

#### 2.1.2 Mathematical principles

SASS identifies hidden and linearly separable data components that are maximally attributable to transcranial electric stimulation and least attributable to brain activity. These components are then rejected to achieve stimulation artifact suppression. SASS is based on a generalized eigenvalue decomposition (joint diagonalization) of encephalographic covariance matrices during and in absence of stimulation. It was shown that using an eigenvalue decomposition of the covariance matrix for Signal-Space Projection (SSP) (Uusitalo & Ilmoniemi, 1997) can be used to suppress tACS artifacts (Vosskuhl et al., 2019), but this approach is agnostic to the spatial distribution of physiological brain activity possibly limiting performance of artifact removal. In SSP, the data matrix becomes linearly separated into orthogonal signal and noise subspaces. This is equivalent to an eigenvalue decomposition of the covariance matrix **A**, where the resulting components *w*_*i*_ (which are orthogonal) can be ordered by the amount of variance *λ*_*i*_ they explain:

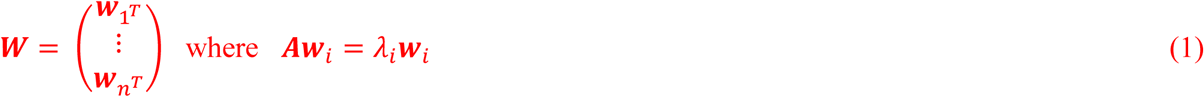

The matrix **P** can then be constructed, which projects the data onto the vector subspace (“signal space”) orthogonal to the top few components explaining the most variance (“noise space”). This can be written as a separation of the data into latent components, followed by a zeroing out of undesired components and a projection back into the original sensor space:

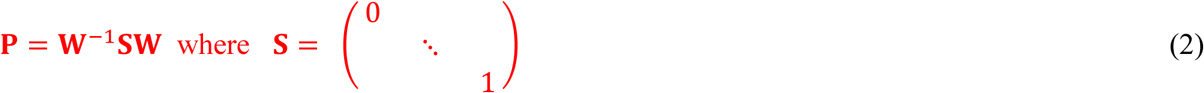

To account for the spatial distribution of physiological brain activity, SASS was designed to identify components that jointly maximize the variance attributable to stimulation artifacts and minimize the variance attributable to brain activity. In SASS, a source separation matrix is computed from the joint diagonalization of EEG sensor covariance matrices during (**A**) and in absence of stimulation (**B**), respectively:

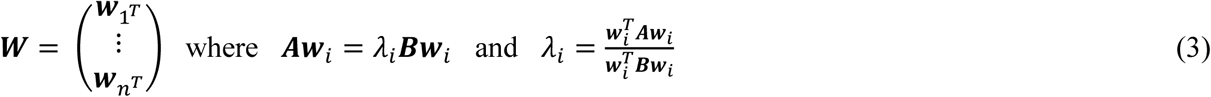

The ratio *λ*_*i*_ represents the ratio of signal power of component *i* during tACS relative to in absence of tACS. Like SSP, SASS can be summarized in a single linear projection:

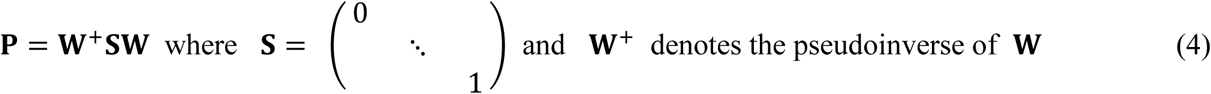

This approach is related to spatio-spectral decomposition (Nikulin, Nolte, & Curio, 2011), which also involves a joint diagonalization of covariance matrices. Like SASS, SSD computes a spatial filter which aims to increase the signal-to-noise ratio of narrowband EEG activity. Different to SASS, SSD contrasts the covariance in the target frequency band against the covariance in neighboring frequency bands. This distinguishes it from SASS, where covariances computed within the same frequency band during and in absence of AM-tACS are contrasted. Importantly, in SASS a certain number of components must be chosen for stimulation artifact rejection. We minimize residual artifacts by selecting the number of components that minimizes the difference in signal power at the target frequency (as measured by mean squared error across all channels (Kasten et al., 2018) between cleaned data recorded during AM-tACS and data recorded in absence of AM-tACS).

### 2.2 Electroencephalography (EEG)

A 64-channel EEG system (Bittium Corp., Oulu, Finland) with passive Ag/AgCl electrodes (Brain Products GmbH, Gilching, Germany) was used to record electrical activity on the scalp. The amplifier was set to DC-mode with a dynamic range of +/−430 V, a resolution of 51 nV/bit, and a range of 24 bit. Electrode impedances were kept below 10 kOhm. Signals were sampled at 500 Hz with an anti-aliasing filter applied at 125 Hz. Saturated electrodes were excluded from the analysis. Electrodes exhibiting broadband power more than two orders of magnitude higher than the median in the condition without AM-tACS were excluded from the analysis.

### 2.3 Transcranial alternating current stimulation (tACS)

AM-tACS was applied to the scalp using a commercial stimulator (NeuroConn GmbH, Ilmenau, Germany). Rubber electrodes with a 4 x 5 cm size were placed over position CPz and on the inion according to the international 10-20 system. This corresponds to the standard montage used for targeting the visual system during tACS experiments (e.g., Helfrich et al., 2014). The stimulator delivered AM-tACS with a 220 Hz carrier and 10 Hz envelope signal. Stimulation intensity was set to a peak-to-peak amplitude of 2 mA.

### 2.4 Presentation of visual flickers

During ongoing AM-tACS, a sinusoidal grating that flickered at 10 Hz was presented for 2 seconds across trials. A random inter-trial interval between 0.5 to 1 second ensured that the onset time of the visual flicker was randomly distributed over the phase of the AM-tACS signal. Visual stimuli were presented via a head-mounted display (Oculus VR Inc., California, USA). Its analog audio output was fed into a bipolar channel of the EEG amplifier and stored to obtain a trigger marker of the stimulus onset time. Jitter between the audio output signal and visual stimulus presentation was under 5 ms. Before the experiment, signal artifacts related to the use of the head-mounted display were ruled out.

### 2.5 Participants and sessions

Seven healthy participants (4 male, 3 female, 22 – 28 years old) were invited to participate in the study and provided written informed consent. The study was approved by the ethics committee of the Charité – University Medicine Berlin (EA1/077/18). Initially, a calibration session consisting of 200 trials of visual flicker was recorded in absence of AM-tACS. Then, a session of 200 trials of visual flickers was recorded while AM-tACS was applied. Recording time for each participant amounted to approximately 20 minutes. One participant was excluded due to a lack of discernible SSVEPs in absence of AM-tACS.

### 2.6 Electroencephalographic (EEG) data processing

MNE-Python (Gramfort et al., 2013) was used for the entire analysis. To obtain a representative channel, unless otherwise specified, analyses were applied to a virtual channel computed from the average of all available occipital electrodes (after SASS, if applied).

#### 2.6.1 Stimulation Artifact Source Separation (SASS)

EEG data in absence and during AM-tACS were bandpass filtered around the stimulation frequency from 9 – 11 Hz using finite impulse response (FIR) filters designed via the Hamming window method (Saramaeki, Mitra, & Kaiser, 1993). The empirical covariance matrix of both unsegmented datasets was then used to compute SASS (Fig. 1 and Section 2.1.2).

#### 2.6.2 Power spectra

To compute power spectra (Fig. 3), Welch periodograms over the entire unfiltered datasets (with or without application of SASS) using 2048 fast Fourier transform (FFT) points were used. To compute high-resolution power spectra (Fig. S5), we followed the procedure of (Noury et al., 2016). We split the entire recordings into 120 second segments and computed Thompson’s multitaper power spectral density (PSD) estimates (0.05 Hz bandwidth, NW=6) for each segment.

**Figure 2:**
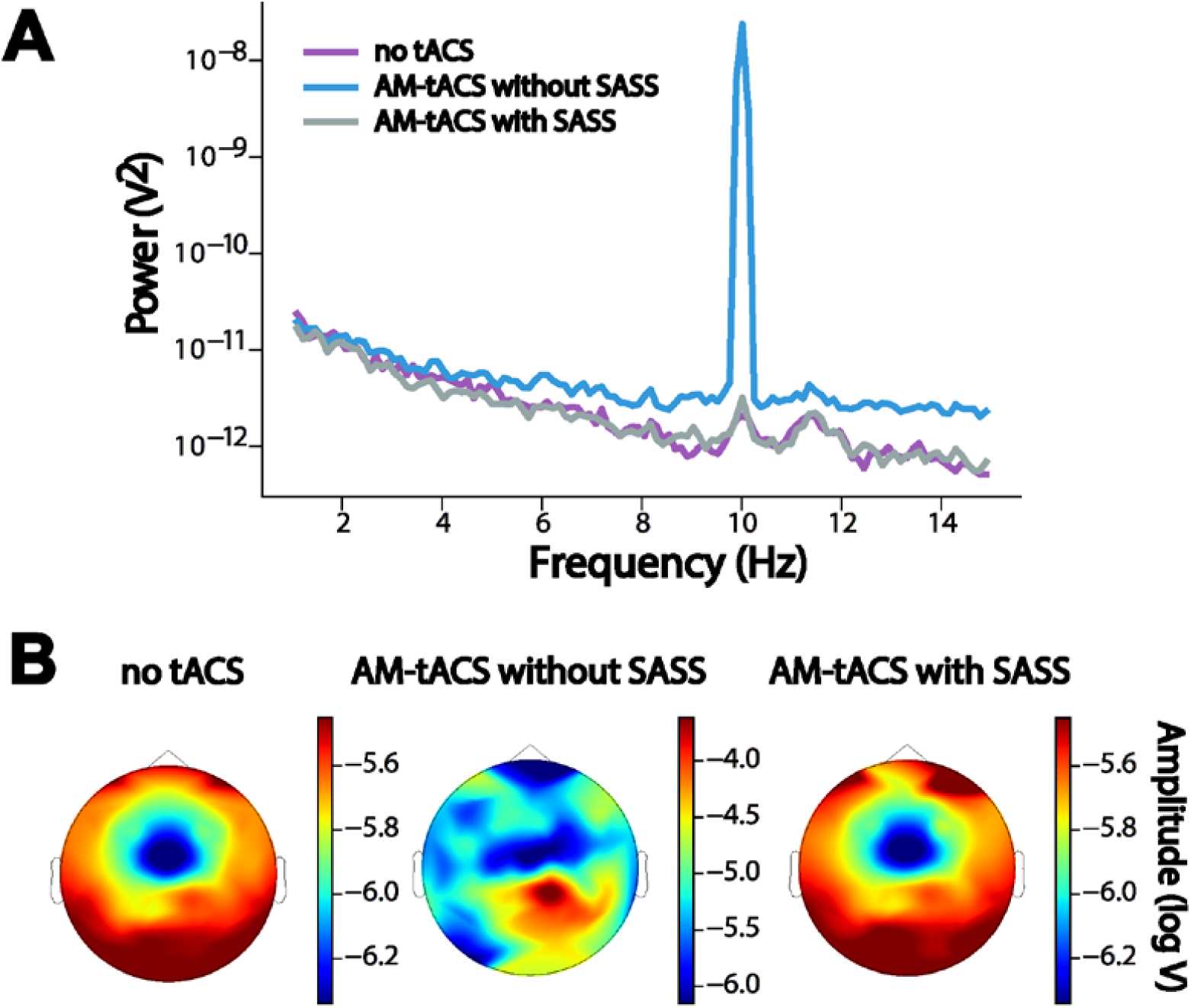
Power spectrum and steady-state visual evoked potential (SSVEP) topography during and in absence of amplitude-modulated transcranial alternating current stimulation (AM-tACS) using Stimulation Artifact Source Separation (SASS) in a representative participant. A: Without SASS, the AM-tACS artifact masked brain activity in the power spectrum at occipital electrodes, including the peak related to SSVEPs at 10 Hz. Using SASS, the physiological power spectrum was recovered. B: Without SASS, the AM-tACS artifact masked the topography of SSVEPs. With SASS, the SSVEP topography was recovered.

**Figure 3:**
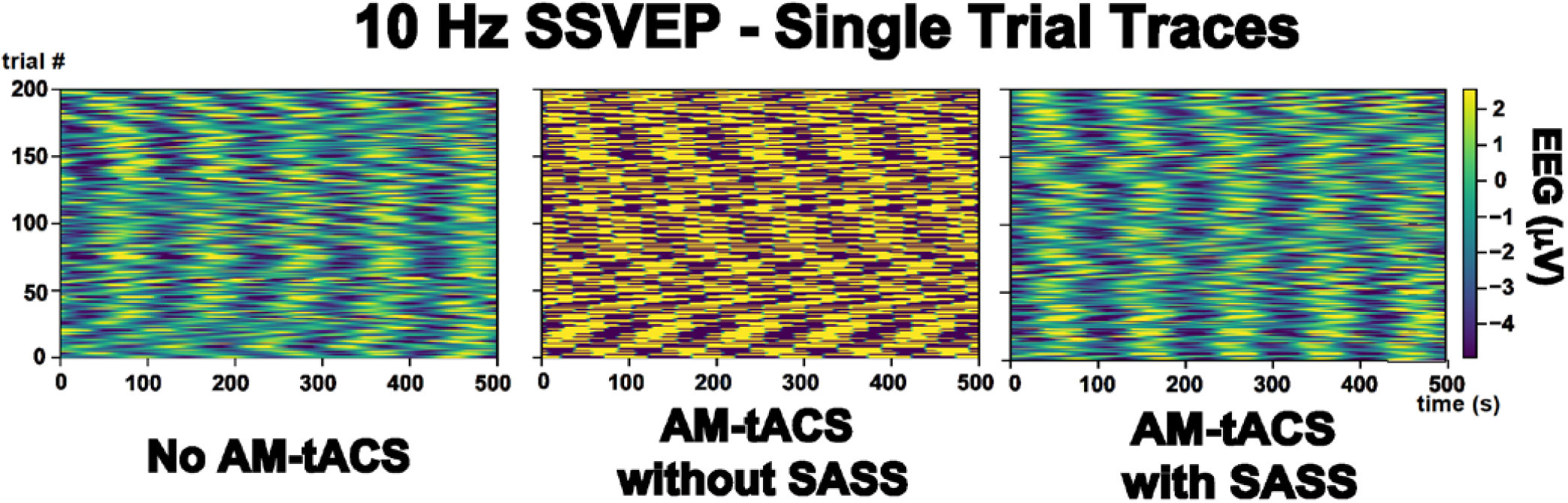
Single-trial steady-state visual evoked potentials (SSVEPs) in absence and during amplitude-modulated transcranial alternating current stimulation (AM-tACS) at the target frequency (10 Hz) with and without using Stimulation Artifact Source Separation (SASS) in a representative participant. In absence of AM-tACS, 10 Hz SSVEPs can be easily identified in single-trial traces of electroencephalographic (EEG) activity (left). When applying AM-tACS targeting the same frequency, i.e. 10 Hz, the AM-tACS artifact masks physiological activity. After applying SASS to EEG data recorded during AM-tACS, physiological single-trial SSVEP activity is recovered.

#### 2.6.3 Modulations of stimulation artifacts by heartbeat

Following (Noury et al., 2016), we assessed modulations of the AM-tACS artifact by heartbeat in the time domain. We FIR filtered the data from 5 – 15 Hz and applied the Hilbert transform to obtain the signal envelope. Then, we windowed the data into 4 second segments centered on electrocardiogram (ECG) R-peaks. We subtracted the temporal mean from each segment and tested for a significant modulation at each timepoint using a permutation test. We compared the average envelope around an R-peak to the average envelopes computed using 1000 random placements of the segments. The resulting p-values were Bonferroni-corrected for multiple comparisons. The permutation test was computed for every channel.

#### 2.6.4 Single-trial amplitude and phase

To compute single-trial amplitude and phase, the unsegmented bandpass filtered EEG data was Hilbert transformed. Then, the data was segmented into 2 second trials corresponding to the presentation of each visual flicker. Finally, instantaneous amplitude and phase difference between EEG and flicker was computed for each timepoint and averaged within each trial to obtain a single data point per trial. For visualization, the mean phase angle across trials was subtracted.

### 2.7 Statistical procedures

To test whether single-trial amplitudes during AM-tACS without SASS were larger than in absence of AM-tACS, we employed a one-sided t-test for independent samples. To test whether single-trial amplitudes during AM-tACS with SASS were smaller than during AM-tACS without SASS, we employed a one-sided t-test for dependent samples. To test whether single-trial amplitudes during AM-tACS with SASS were different from single-trial amplitudes in absence of AM-tACS (i.e. whether any residual artifacts remained after SASS), we employed a two-sided t-test for independent samples. A power analysis revealed that an effect size of 0.281 (Cohen’s d) could be detected with 0.8 power at the employed alpha level of 0.05, meaning that a difference in means (residual artifact) of between 0.174 and 0.972 μV (depending on the standard deviations of single-trial amplitudes within a subject) could be detected.

Analogously to the case of single-trial amplitudes, we employed Wallraff tests (Zar, 1999) for dependent or independent samples to test for differences in single-trial phases (relative to the visual flicker) within each subject. In the dependent samples case, angular distances were compared using a Wilcoxon rank-sum test. In the independent samples case, angular distances were compared using a Mann-Whitney U test. Phase locking values and mean single-trial amplitudes at the group level were statistically compared using one- or two-sided Wilcoxon rank-sum tests.

To test for differences in single-trial amplitude between conditions across the entire sensor space, spatial cluster-based permutation tests with threshold-free cluster enhancement (Smith & Nichols, 2009) were computed individually for each participant. For each sensor and participant, 200 data points corresponding to the SSVEP trials were used.

## 3. Results

### 3.1 SASS recovered the power spectrum and topography of electric brain oscillations

Fig. 2 depicts successful recovery of the occipital power spectrum and SSVEP amplitude topography during AM-tACS using SASS. Of note, a clear peak in the power spectrum at 10 Hz remained due to steady-state visual stimuli presented at 10 Hz. To test for differences in the spatial distribution of SSVEP amplitude between EEG recorded in absence of AM-tACS and EEG recorded during AM-tACS, spatial cluster-based permutation tests were performed individually for each participant. No significant difference (i.e., residual artifact) was detected for any of the participants.

### 3.2 SASS recovered single-trial amplitude and phase of electric brain oscillations

Fig. 3 depicts recovery of individual SSVEP traces by SASS. Single-trial SSVEP amplitude was significantly increased (by multiple orders of magnitude) during AM-tACS when compared to SSVEP amplitudes recorded in absence of AM-tACS. Applying SASS, across all participants, SSVEP amplitudes during AM-tACS was comparable to SSVEP amplitudes recorded in absence of AM-tACS (Fig. 4A, Table 1). A power analysis revealed that, using SASS, a difference in means (residual artifact) of above 0.174 and 0.972 μV, respectively (depending on the variances of single-trial amplitudes within a subject), could have been detected (see Section 2.7). Single-trial SSVEP phase (relative to the visual flicker) was significantly distorted during AM-tACS compared to data recorded in absence of AM-tACS. Using SASS, single-trial SSVEP phase during AM-tACS was not different from single-trial SSVEP phase recorded in absence of AM-tACS (Fig. 4B, Table 2). Figs. S3 and S4 depict the results seen in Fig. 4 for all study participants.

**Table 1.** Single-trial amplitude of steady-state visual evoked potentials (SSVEPs) across all participants. Two-sided t-tests (for dependent or independent samples, as applicable) were used for pairwise comparisons of single-trial amplitudes between conditions. Mean (and standard deviations) are reported.

| Participant | EEG amplitude in absence of tACS (μV) | EEG amplitude during tACS without SASS (μV) | EEG amplitude during tACS with SASS (μV) | P-value (absence of tACS vs. tACS without SASS) | P-value (tACS without SASS vs. tACS with SASS) | P-value (absence of tACS vs. tACS with SASS) |
| --- | --- | --- | --- | --- | --- | --- |
| 1 | 3.13 (0.652) | 27.3 (1.21) | 3.19 (0.689) | <0.001 | <0.001 | 0.396 |
| 2 | 7.70 (3.47) | 81.9 (30.0) | 8.20 (3.45) | <0.001 | <0.001 | 0.149 |
| 3 | 2.85 (0.980) | 89.4 (2.82) | 2.90 (0.686) | <0.001 | <0.001 | 0.605 |
| 4 | 5.60 (1.53) | 799 (211) | 5.57 (1.31) | <0.001 | <0.001 | 0.832 |
| 5 | 5.73 (2.28) | 13.2 (3.56) | 5.92 (1.56) | <0.001 | <0.001 | 0.371 |
| 6 | 2.38 (0.679) | 23.2 (0.871) | 23.1 (0.562) | <0.001 | <0.001 | 0.309 |

**Table 2.** Single-trial phase of steady-state visual evoked potentials (SSVEPs) across all participants. Wallraff tests (for dependent or independent samples, as applicable) were used for pairwise comparisons of the distribution of SSVEP-flicker phase differences between conditions. Phase locking value (PLV) is reported.

| Participant | Phase locking in absence of tACS (PLV) | Phase locking during tACS without SASS (PLV) | Phase locking during tACS with SASS (PLV) | P-value (absence of tACS vs tACS without SASS) | P-value (tACS without SASS vs tACS with SASS) | P-value (absence of tACS vs tACS with SASS) |
| --- | --- | --- | --- | --- | --- | --- |
| 1 | 0.457 | 0.0437 | 0.350 | <0.001 | <0.001 | 0.0974 |
| 2 | 0.181 | 0.046 | 0.217 | <0.05 | <0.05 | 0.533 |
| 3 | 0.536 | 0.0131 | 0.412 | <0.001 | <0.001 | 0.160 |
| 4 | 0.531 | 0.0355 | 0.531 | <0.001 | <0.001 | 0.609 |
| 5 | 0.479 | 0.0900 | 0.371 | <0.001 | <0.001 | 0.346 |
| 6 | 0.268 | 0.0458 | 0.252 | <0.01 | <0.05 | 0.936 |

**Figure 4:**
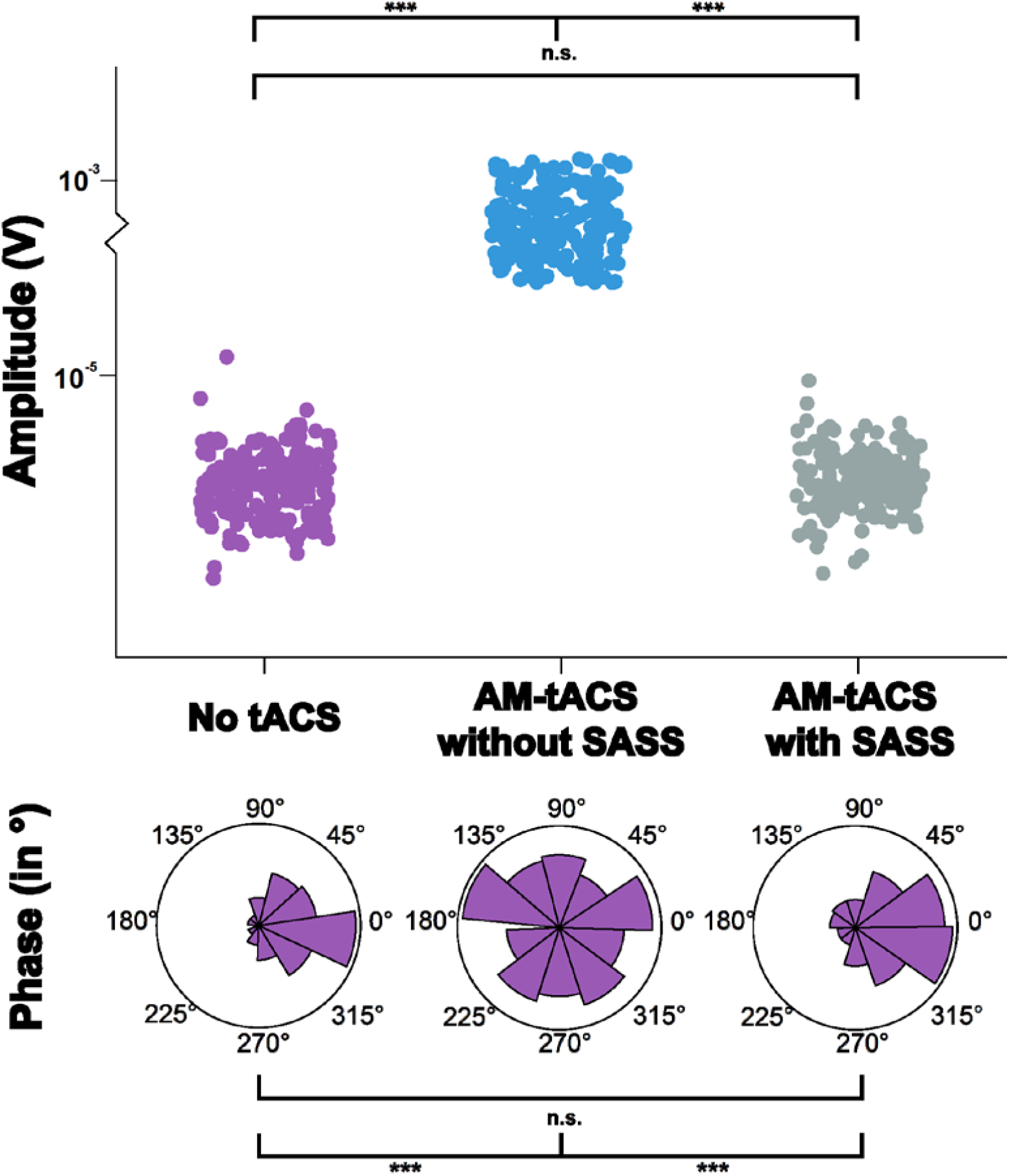
Single-trial amplitude (A) and phase (B) of steady-state visual evoked potentials (SSVEPs) during and in absence of amplitude-modulated transcranial alternating current stimulation (AM-tACS) using Stimulation Artifact Source Separation (SASS) in a representative participant. A: Single-trial amplitude of electroencephalographic (EEG) data at 10 Hz recorded in absence and during AM-tACS using SASS. When applying SASS, single-trial amplitudes were comparable to activity recorded in absence of AM-tACS. B: Single-trial phase of EEG data at 10 Hz recorded in absence and during AM-tACS relative to the phase of the visual flicker. When applying SASS, similar to absence of AM-tACS, single-trial phase was locked to the visual flicker during AM-tACS.

### 3.3 Stimulation Artifact Source Separation (SASS) recovered mean amplitude and phase locking value (PLV) of electric brain oscillations at the group level

Fig. 5 depicts summary statistics of the performance of SASS at group level. Mean amplitudes of 10 Hz activity during AM-tACS (172 ± 282 μV) were significantly higher compared EEG data recorded in absence of AM-tACS (4.57 ± 1.92 μV). We found no difference in 10 Hz activity in data recorded in absence and during AM-tACS when applying SASS (4.68 ± 2.07 μV). Phase locking between EEG data and visual flickers (Fig. 5B) decreased during AM-tACS (0.0456 ± 0.0228 PLV) compared to EEG data recorded in absence of AM-tACS (0.409 ± 0.135 PLV). When applying SASS, phase locking between EEG data and visual flickers was not different during AM-tACS compared to EEG data recorded in absence of stimulation (0.355 ± 0.103 PLV).

**Figure 5:**
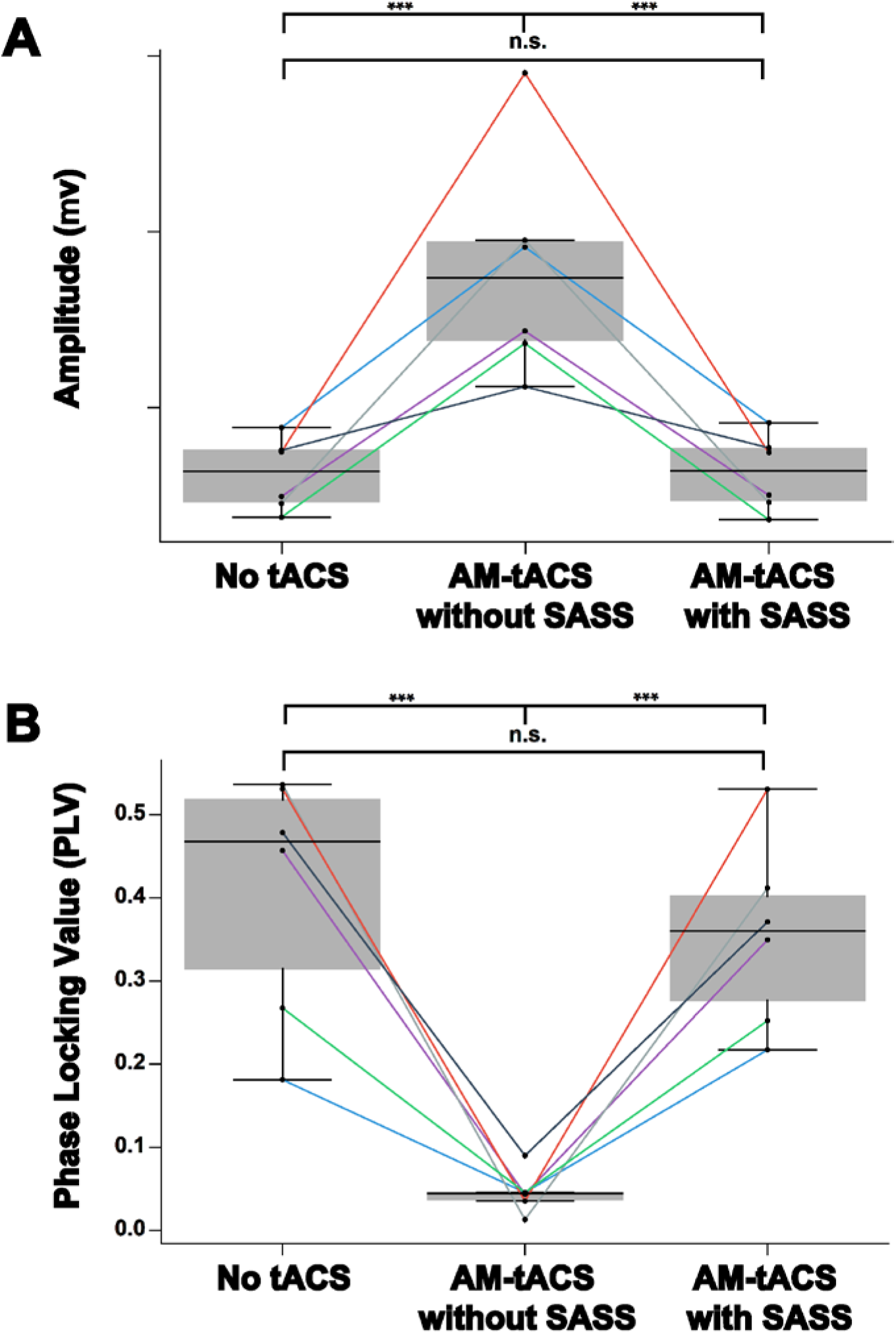
Restoration of amplitude and phase of steady-state visual evoked potentials (SSVEPs) using Stimulation Artifact Source Separation (SASS) at group level. A: When not applying SASS, mean single-trial amplitudes of 10 Hz electroencephalographic (EEG) activity increased indicating artifact distortions. When applying SASS, mean single-trial 10 Hz EEG amplitudes during amplitude-modulated transcranial alternating current stimulation (AM-tACS) were comparable to EEG data recorded in absence of stimulation. B: When not applying SASS, phase locking of 10 Hz EEG activity to the visual flicker decreased indicating artifact distortions of SSVEP. When applying SASS, phase locking of 10 Hz EEG activity to the visual flicker was recovered. Data from occipital sensors of all six study participants is depicted.

### 3.4 Properties of Stimulation Artifact Source Separation (SASS)

Depending on the experimental design and purpose of study, there are a number of properties of SASS that users should consider. First, presumably due to spatially varying capacitive effects (Noury & Siegel, 2017a), the topography of the AM-tACS artifact is frequency-dependent. Therefore, SASS must be computed separately for each frequency band of interest (Fig. S7). Second, the optimal number of components to reject (Section 2.1.2) may vary across participants. The eigenvalue spectrum and spatial patterns of rejected components across all subjects are visualized in Fig. S6. Third, applying a time- and frequency-domain analysis for a selected occipital channel across all study participants according to (Noury et al., 2016) did not evidence any demodulations including intermodulation distortions at the AM-tACS target frequency (Fig. S5). Fourth, due to time-varying covariances, SASS computed for one dataset should not be applied to another dataset. This may lead to sub-optimal performance (Fig. S8). Therefore, we advise to recompute SASS according to the covariance of the dataset it is applied to. This is particularly true for online applications, where recursive solutions to the underlying eigenvalue problem could be applied (Li, Yue, Valle-Cervantes, & Qin, 2000). Finally, SASS does not overcorrect the signals of interest when computed as usual but applied to the dataset in absence of AM-tACS. We have used a spatial cluster-based permutation test (Section 2.7) to verify that single-trial amplitudes are not attenuated in any of the study participants due to the application of SASS to data recorded in absence of AM-tACS.

## 4. Discussion

Up to now, there was no real-time compatible signal processing tool available for AM-tACS artifact suppression that allows for recovering phase and amplitude of evoked brain responses at a single-trial level. Here, we introduced SASS, a spatial filter based on a source separation matrix computed from joint diagonalization of EEG sensor covariance matrixes at the target frequency recorded in absence and during AM-tACS. To evaluate the effectiveness of SASS, we used a 10-Hz-SSVEP paradigm resulting in evoked oscillatory brain responses with known frequency, amplitude and phase. Electric brain activity was assessed using EEG across six healthy volunteers in absence and during AM-tACS targeting the SSVEP frequency. Using SASS, single-trial SSVEP amplitude and phase information was restored to the level recorded in absence of stimulation (Fig. 4) across all study participants. Likewise, the SSVEP topography was successfully recovered (Fig. 3) across all study participants.

Besides showing that SASS is an effective tool for separating electric brain activity and stimulation artifacts at the frequency targeted by AM-tACS, our results pave the way for implementation of adaptive stimulation paradigms in which the phase, amplitude and spatial distribution of the applied electric field is adapted to ongoing brain oscillations. This may contribute to the development of more effective stimulation approaches to target various brain functions and further elucidate the underlying mechanisms of AM-tACS.

While 200 trials employed in the current study provide sufficient statistical power to detect residual artifacts of above 0.174 and 0.972 μV, respectively, (Section 2.7) averaging of instantaneous phase differences and amplitudes within each single trial may have masked residual artifacts. Furthermore, in SASS, the sensor covariance matrix of physiological signals of interest is estimated from EEG data recorded in absence of stimulation. However, this covariance may be affected by AM-tACS via modulations of brain activity. Therefore, it cannot be ruled out that signal of interests are also attenuated when applying SASS during AM-tACS (e.g., entrained endogenous brain oscillations). Further studies are required to systematically assess this possibility. A similar issue may occur when SASS is precomputed and applied to novel data (Fig. S8). Due to time-varying covariances, robust online application of SASS will require methods for dynamically solving the eigenvalue problem and updating the spatial filter. This can be achieved by a recursive solution to the underlying eigenvalue problem, commonly used in real-time systems in other fields (Li et al., 2000). Future work will also need to include a comprehensive validation for other electrode montages, exploring how the spatial relationship of cortical sources and stimulation electrodes affects the performance of SASS. Finally, while in this work we strived to validate SASS for AM-tACS, application of SASS in combination with other stimulation protocols, e.g. conventional tACS, is also possible. However, depending on the magnitude of artifacts at the frequencies of interest, more signal components may have to be rejected reducing the dimensionality of data.

Despite these considerations, we have shown that SASS allows for successful recovery of single-trial SSVEP amplitude and phase during AM-tACS. Thereby, SASS now allows for further investigation of tACS-related network effects (Alekseichuk et al., 2019; Reinhart & Nguyen, 2019). SASS could also be used to purposefully modulate large-scale synchronization (Reinhart & Nguyen, 2019) by targeting one cortical region as a function of another. Moreover, SASS may also help to better understand the underlying mechanisms of tACS effects. Recent studies suggest that tACS effects are not only mediated by electric field-dependent modulations of membrane potentials in superficial cortical layers, but may also involve transcutaneous stimulation of skin nerves (Asamoah, Khatoun, & Mc Laughlin, 2019). Here, SASS may help to identify the primary mechanism of action based on precise characterization of phase locking and phase lags in combination with other neurophysiological measures such as neural conduction times and cortico-cortical phase synchronization.

Real-time phase estimation of ongoing brain oscillations is challenging and critically depends on the signal-to-noise ratio and instantaneous amplitude of the signal (Zrenner et al., 2020). It is thus important to note that successful implementation of adaptive AM-tACS not only requires real-time suppression of stimulation artifacts, but also stable phase estimation accuracy. This could be achieved by purposefully amplifying the target oscillation’s amplitude using a cognitive task (e.g. motor imagery to increase μ-rhythm amplitude) (Soekadar, Witkowski, Birbaumer, & Cohen, 2015; Soekadar, Witkowski, Cossio, Birbaumer, & Cohen, 2014), sensory stimuli (e.g. visual flickers or vibrotactile stimulation), or operant conditioning (e.g. neurofeedback) (Ruddy et al., 2018).

## Data availability

The data is publicly available on Mendeley Data: https://data.mendeley.com/datasets/39n9zttp4t. The implementation of the novel algorithm (Stimulation Artifact Source Separation, SASS) is publicly available on GitHub: https://github.com/davidhaslacher/sass.

## Disclosures

None.

## CRediT authorship contribution statement

D.H., K.N., S.R.S., C.B. and S.E.R. were involved in the design of study, analysis and/or interpretation of data, drafting the manuscript, and revising the manuscript critically for important intellectual content. D.H. and K.N. acquired the data.

## Acknowledgements

This work was supported in part by the European Research Council (ERC) under the project NGBMI (759370), the Baden-Württemberg Stiftung (NEU007/1) and Einstein Stiftung Berlin. SRS received special support by the Brain & Behavior Research Foundation as 2017 NARSAD Young Investigator Grant recipient and P&S Fund Investigator.

## Supplementary Materials (prepared for submission as ‘Data in Brief’)

**Fig. S1. Power spectrum of occipital sensors across all participants.**

**Fig. S2. Topography of mean single-trial steady-state visual evoked potentials (SSVEPs) amplitude across all participants.**

**Fig. S3. Amplitude of single-trial steady-state visual evoked potentials (SSVEPs) across all participants.**

**Fig. S4. Phase of single-trial steady-state visual evoked potentials (SSVEPs) (relative to flicker) across all participants.**

**Fig. S5. Assessment of possible nonlinear modulations of the amplitude-modulated transcranial alternating current stimulation (AM-tACS) artifact at the target frequency (10 Hz), e.g. by physiological activity (heartbeats).** After applying time- and frequency-domain analyses according to (Noury et al., 2016; Noury & Siegel, 2017b) around the AM-tACS target frequency (10 Hz) at channel O2, no artifacts neighboring the frequency of interest in the power spectrum or time domain could be detected (see Section 2.6).

**Fig. S6. Eigenvalue spectrum (a) and spatial patterns of first six rejected components (b) for each study participant.** The generalized eigenvalue is depicted above each component.

**Fig. S7. Topography of amplitude-modulated transcranial alternating current stimulation (AM-tACS) artifacts is frequency dependent.** AM-tACS artifacts, like tACS artifacts, appear at the stimulation frequency and its harmonics. However, the topography of the artifact is frequency-dependent (bottom), presumably due to spatially varying nonlinear transformations of the current by different capacitive effects at each electrode (Noury & Siegel, 2017b). Therefore, Stimulation Artifact Source Separation (SASS) should be computed separately for each frequency of interest.

**Fig. S8. Performance of Stimulation Artifact Source Separation (SASS) on novel data.**

SASS was computed on the first half of each AM-tACS dataset and tested on the second half.

## Notes

### Competing Interest Statement

The authors have declared no competing interest.

